# Brain Atlas for Glycoprotein Hormone Receptors at Single-Transcript Level

**DOI:** 10.1101/2022.06.01.494351

**Authors:** Vitaly Ryu, Anisa Gumerova, Funda Korkmaz, Seong Su Kang, Pavel Katsel, Sari Miyashita, Hasni Kannangara, Liam Cullen, Pokman Chan, Tanchun Kuo, Ashley Padilla, Samir Zaidi, Se-Min Kim, Maria I. New, Clifford J. Rosen, Ki A. Goosens, Tal Frolinger, Vahram Haroutunian, Keqiang Ye, Daria Lizneva, Terry F. Davies, Tony Yuen, Mone Zaidi

**Affiliations:** Center for Translational Medicine and Pharmacology, Icahn School of Medicine at Mount Sinai, New York, NY 10029; Department of Medicine and of Pharmacological Sciences, Icahn School of Medicine at Mount Sinai, New York, NY 10029; Department of Pathology, Emory University School of Medicine, Atlanta, GA 30322; Department of Psychiatry, Icahn School of Medicine at Mount Sinai, New York, NY 10029; Alamak Biosciences, Beverly, MA 01915; Memorial Sloan Kettering Cancer Center, New York, NY 10065; Department of Pediatrics, Icahn School of Medicine at Mount Sinai, New York, NY 10029; Maine Medical Center Research Institute, Scarborough, ME 04074; Faculty of Life and Health Sciences, and Brain Cognition and Brain Disease Institute, Shenzhen Institute of Advanced technology, Chinese Academy of Sciences, Shenzhen, China

## Abstract

There is increasing evidence that anterior pituitary hormones, traditionally thought to have unitary functions in regulating single endocrine targets, act on multiple somatic tissues, such as bone, fat, and liver. There is also emerging evidence for anterior pituitary hormone action on brain receptors in mediating central neural and peripheral somatic functions. Here, we have created the most comprehensive neuroanatomical atlas on the expression of TSHRs, LHCGRs and FSHRs. We have used RNAscope, a technology that allows the detection of mRNA at single-transcript level, together with protein level validation, to document *Tshr* expression in 173 and *Fshr* expression in 353 brain regions, nuclei and sub–nuclei identified using the *Atlas for the Mouse Brain in Stereotaxic Coordinates*. We also identified *Lhcgr* transcripts in 401 brain regions, nuclei and sub–nuclei. Complementarily, we used ViewRNA, another single-transcript detection technology, to establish the expression of *FSHRs* in human brain samples, where transcripts were co–localized in *MALAT1–positive* neurons. In addition, we show high expression for all three receptors in the ventricular region—with yet unknown functions. Intriguingly, *Tshr* and *Fshr* expression in the ependymal layer of the third ventricle was similar to that of the thyroid follicular cells and testicular Sertoli cells, respectively. TSHRs were expressed specifically in tanycytes. In contrast, *Fshrs* were localized to NeuN–positive neurons in the granular layer of the dentate gyrus in murine and human brain—both are Alzheimer’s disease vulnerable regions. Our atlas thus provides a vital resource for scientists to explore the link between the stimulation or inactivation of brain glycoprotein hormone receptors on somatic function. New actionable pathways for human disease may be unmasked through further studies.

## INTRODUCTION

There is increasing evidence that pituitary hormones traditionally thought of as ‘pure’ regulators of single physiological processes affect multiple bodily systems, either directly or *via* actions on brain receptors (1, 2). We established, for the first time, a direct action of thyroid–stimulating hormone (TSH) on bone and found that TSH receptor (TSHR) haploinsufficiency causes profound bone loss in mice (2). We also found that follicle–stimulating hormone (FSH), *hitherto* thought to solely regulate gonadal function, displayed direct effects on the skeleton to cause bone loss (3), and on fat cells, to cause adipogenesis and body fat accumulation (4). Likewise, we showed that hormones from the posterior pituitary, namely oxytocin and vasopressin, displayed direct, but opposing skeletal actions—effects that may relate to the pathogenesis of bone loss in pregnancy and lactation, and in chronic hyponatremia, respectively (5–8). To add to this complexity, and in addition to the poorly recognized ubiquity of pituitary hormone receptors, the ligands themselves, or their variants, are expressed widely. We find the expression of a TSHβ variant (TSHβv) in bone marrow macrophages, while oxytocin is expressed by both osteoblasts and osteoclasts (9–12). These studies have together shifted the paradigm from established unitary functions of pituitary hormones to an evolving array of yet unrecognized roles of physiologic and pathophysiologic importance.

There is a compelling body of literature to support the expression of oxytocin receptors in various brain regions, and their function in regulating peripheral actions, such as social behavior and satiety (5, 13). However, there is relatively scant information on the expression, and importantly, the function of the anterior pituitary glycoprotein hormone family of receptors, namely FSHR, TSHR and luteinizing hormone/human chorionic gonadotropin receptor (LHCGR). Discrete sites of the rat, mouse and human brain express receptors for these hormones, with several studies pointing to their relationship to neural functions, such as cognition, learning, neuronal plasticity, and sensory perception, as well as to neuropsychiatric disorders, including affective disorders and neurodegeneration (14–21) (Table 1). In the light of such discoveries, the link between the stimulation of these receptors in the brain and the regulation of peripheral physiological processes needs further investigation.

**Table 1:** Known Functions of TSHR, FSHR and LHCGR in Brain.

| Receptor | Species | Brain Region | Possible Function | Reference |
| --- | --- | --- | --- | --- |
| TSHR | Rat | Hypothalamus | Aging | (15) |
|  | Mice | Hippocampus | Spatial learning and memory | (17) |
|  | Rat | Hypothalamus, hippocampus, pyriform and postcingulate cortex | Thyroid regulation | (14) |
|  | Rat | Hypothalamus | Feeding behavior | (57) |
|  | Human | Hypothalamus, amygdala, cingulate gyrus, frontal cortex, hippocampus, thalamus | Mood disorders | (21) |
|  | Quail | Hypothalamus | Seasonal reproduction | (32) |
| FSHR | Yak | Hypothalamus, pineal gland | Follicle growth, maturation, estrus | (58) |
|  | Mice | Hippocampus | Mood regulation | (18) |
| LHCGR | Rat | Hypothalamus | Aging | (15) |
|  | Mice | Hippocampus, cortex | Spatial memory, cognition, plasticity | (19) |
|  | Rat | Hippocampus | Brain metabolism | (27) |
|  | Fish | Hypothalamus | Functional roles | (59) |
| | Mice | Hippocampus | Promote Amyloid-β formation | (60) |
|  | Mice | Cortex | Regulation of neurosteroid production | (20) |
|  | Mice | Hypothalamus, hippocampus, midbrain, cortex | Regulation of reproductive functions | (61) |
|  | Yak | Hypothalamus, pineal gland | Follicle growth, maturation, estrus | (58) |
|  | Rat | Hypothalamus, hippocampus, dentate gyrus, cerebellum, brainstem, cortex | Cognitive function (AD) | (16) |

Here, we use RNAscope—a cutting–edge technology that detects single RNA transcripts—to create the most comprehensive atlas of glycoprotein hormone receptors in mouse brain. This compendium of glycoprotein hormone receptors in concrete brain regions and sub–regions at a single-transcript level should allow investigators to study both peripheral and central effects of the activation of individual receptors in health and disease. Our identification of brain nuclei with the highest density for each receptor should also create a new way forward in understanding the functional engagement of receptor–bearing nuclei within a large–scale functional network.

## RESULTS

Very little is known about the function(s) of anterior pituitary hormone receptors in the brain, except for isolated studies showing a relationship with cognition and affect (Table 1). We therefore used RNAscope to map the expression of *Tshr, Lhcgr* and *Fshr* in the mouse brain; immunofluorescence and qPCR to provide confirmatory evidence for *Tshr* and *Fshr* expression; and ViewRNA and qPCR to examine for *FSHR* expression in AD–vulnerable regions of the human brain. RNAscope, which allows the detection of single transcripts, uses ~20 pairs of transcript–specific double *Z*–probes to hybridize 10–μm–thick whole brain sections. Preamplifiers first hybridize to the ~28–bp binding site formed by each double *Z*–probe; amplifiers then bind to the multiple binding sites on each preamplifier; and finally, labeled probes containing a fluorescent molecule bind to multiple sites of each amplifier. RNAscope data was quantified on sections from coded mice. Each section was viewed and analyzed using CaseViewer 2.4 (3DHISTECH, Budapest, Hungary) or QuPath v.0.2.3 (University of Edinburgh, UK). The *Atlas for the Mouse Brain in Stereotaxic Coordinates* (22) was used to identify every nucleus or sub–nucleus in which we manually counted *Tshr, Lhcgr* or *Fshr* transcripts in every tenth section using a tag feature. Repeat counting of the same section agreed within <2%. Receptor density was calculated by dividing transcript count by the total area (μm^2^, ImageJ) of each region, nucleus or sub–nucleus. Photomicrographs were prepared using Photoshop CS5.1 (Adobe) only to adjust brightness, contrast and sharpness, to remove artifacts (i.e., obscuring bubbles), and to make composite plates.

*Tshrs* were detected bilaterally in 173 brain nuclei and sub–nuclei, in the following descending order of transcript densities: ventricular region, olfactory bulb, forebrain, hypothalamus, medulla, cerebellum, midbrain and pons, cerebral cortex, hippocampus and thalamus (Fig. 1A). Importantly, thyroid glands from *Tshr^−/−^* mice did not show a signal, proving probe specificity (Fig. 1B). *Tshr* expression in pooled brain samples was confirmed by qPCR (Fig. 1C). The hypothalamus and hippocampus expressed *Tshr* s, with hypothalamic expression being considerably higher (*P*<0.01) in females than in males. Furthermore, within other regions of the brain, highest *Tshr* densities were as follows: ependymal layer of the third ventricle (slightly higher than the thyroid follicular cells); VTT in the olfactory bulb; HDB in the forebrain; MTu in the hypothalamus; SoIV in the medulla; PFI in the cerebellum; LDTg in midbrain and pons; DP in the cerebral cortex; DG in hippocampus; and PPT in the thalamus (Fig. 1D) (see Appendix for nomenclature). Raw transcript counts in each region and representative micrographs are shown in Supplementary Figs. 1 and 2, respectively.

**Figure 1:**
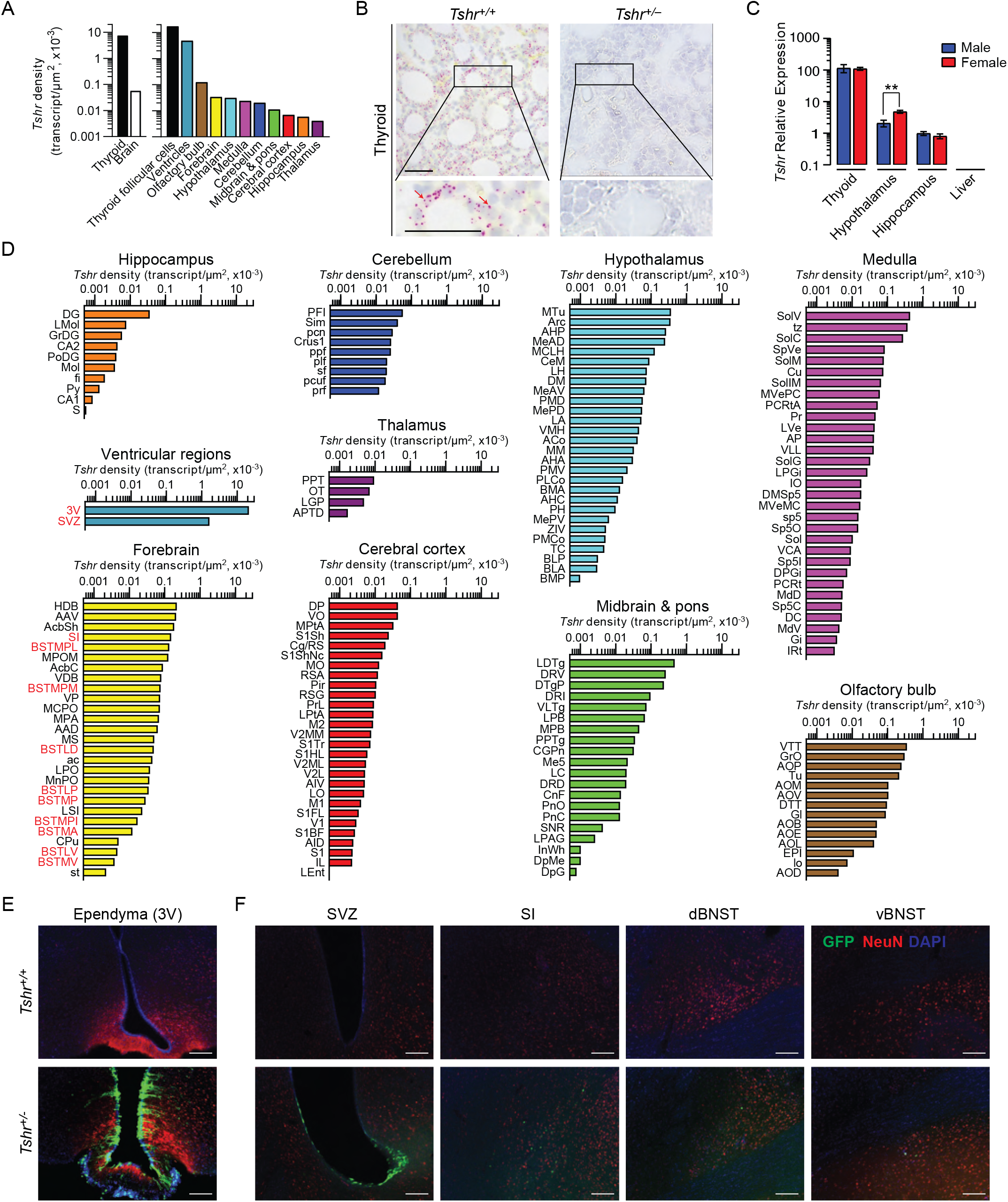
*Tshr* Expression in the Mouse Brain. **(A)** *Tshr* transcript density in the thyroid and various brain regions detected by RNAscope. **(B)** RNAscope probe specificity is confirmed in the *Tshr^+/+^* thyroid. *Tshr^-/-^* thyroid was used as negative control. Scale bar: 50 μm. **(C)** *Tshr* expression in the mouse hypothalamus and hippocampus using quantitative PCR. The thyroid and the liver serve as positive and negative controls, respectively. Statistics: Mean ± s.e.m., *N*=4–5 mice/group, \*\**P*<0.01. **(D)** *Tshr* transcript density in nuclei and sub-nuclei of the ventricular regions, olfactory bulb, forebrain, hypothalamus, medulla, cerebellum, midbrain & pons, cerebral cortex, hippocampus and thalamus. **(E)** Abundant GFP immunofluorescence (green) was detected in the ependymal layer of the third ventricle in *Tshr^+/-^* heterozygous mice, in which a GFP cassette replaced exon 1 of the *Tshr* gene. This GFP signal was absent in *Tshr^+/+^* mice. **(F)** GFP immunofluorescence was also detected in the subventricular zone (SVZ) of the lateral ventricle, and substantia innominata (SI) and dorsal and ventral bed nucleus of stria terminalis (BNST) in the forebrain of the *Tshr^+/-^* mice. Sections were co-stained with DAPI (blue) and a neuronal marker, NeuN (red). Scale bar: 100 μm.

For purposes of replicability, we employed a complementary approach to study brain *Tshr* expression—the *Tshr*–deficient mouse—in which exon 1 of the *Tshr* gene is replaced by a *Gfp* cassette. This reporter strategy allows for the *in vivo* display of *Tshr* locations using GFP immunoreactivity (GFP–ir) as a surrogate for *Tshr* expression (2). Of note is that the *Tshr^+/−^* (haploinsufficient) mouse has one *Tshr* allele intact with normal thyroid function but expresses GFP *in lieu* of one lost allele. In contrast, the *Tshr^+/+^* mouse does not express GFP-ir because both *Tshr* copies are intact and is therefore our negative control.

Consistent with our RNAscope finding, profound GFP–ir was noted in the ependymal region of the third ventricle, mostly in NeuN–negative tanycytes, but with some neuronal localization (Fig. 1E). The SVZ of the lateral ventricles, and the SI, and dorsal and ventral BNST of the forebrain also showed GFP-ir, but immunoreactivity was much lower than the ependymal layer of the third ventricle (Fig. 1F). In all, while there was overall concordance between the two methodologies for high *Tshr*–expressing areas, GFP–ir was not detected in a number of *Tshr*–positive regions. This latter discrepancy most likely reflects the grossly lower sensitivity of immunohistochemical detection.

There is evidence that high LH levels in post–menopausal women correlate with a higher incidence of Alzheimer’s disease (AD) (23, 24); that LHβ transgenic mice are cognitively impaired (25); that LH receptors (LHCGRs) are present in the hippocampus (26, 27); and that hCG induces cognitive deficits in rodents (28, 29). Thus, we mapped *Lhcgr* s in mouse brain to document expression in 401 brain nuclei and sub–nuclei. Probe specificity was established by a positive signal in testicular Leydig cells, and with an absent signal in juxtaposed Sertoli cells (Fig. 2A). Notably similar to *Tshr* transcripts, the ventricular regions displayed the highest transcript density (Fig. 2B). Among the brain divisions, the densities were as follows: OV in the ventricular region; SFO in the forebrain; PFI in the cerebellum; MiA in the olfactory bulb; SCO in the thalamus; PMD in the hypothalamus; MVPO in the medulla; DT in midbrain and pons; GrDG in the hippocampus; and SL in the cerebral cortex (Fig. 2C). Raw transcript counts in each region and representative micrographs are shown in Supplementary Figs. 3 and 4, respectively.

**Figure 2:**
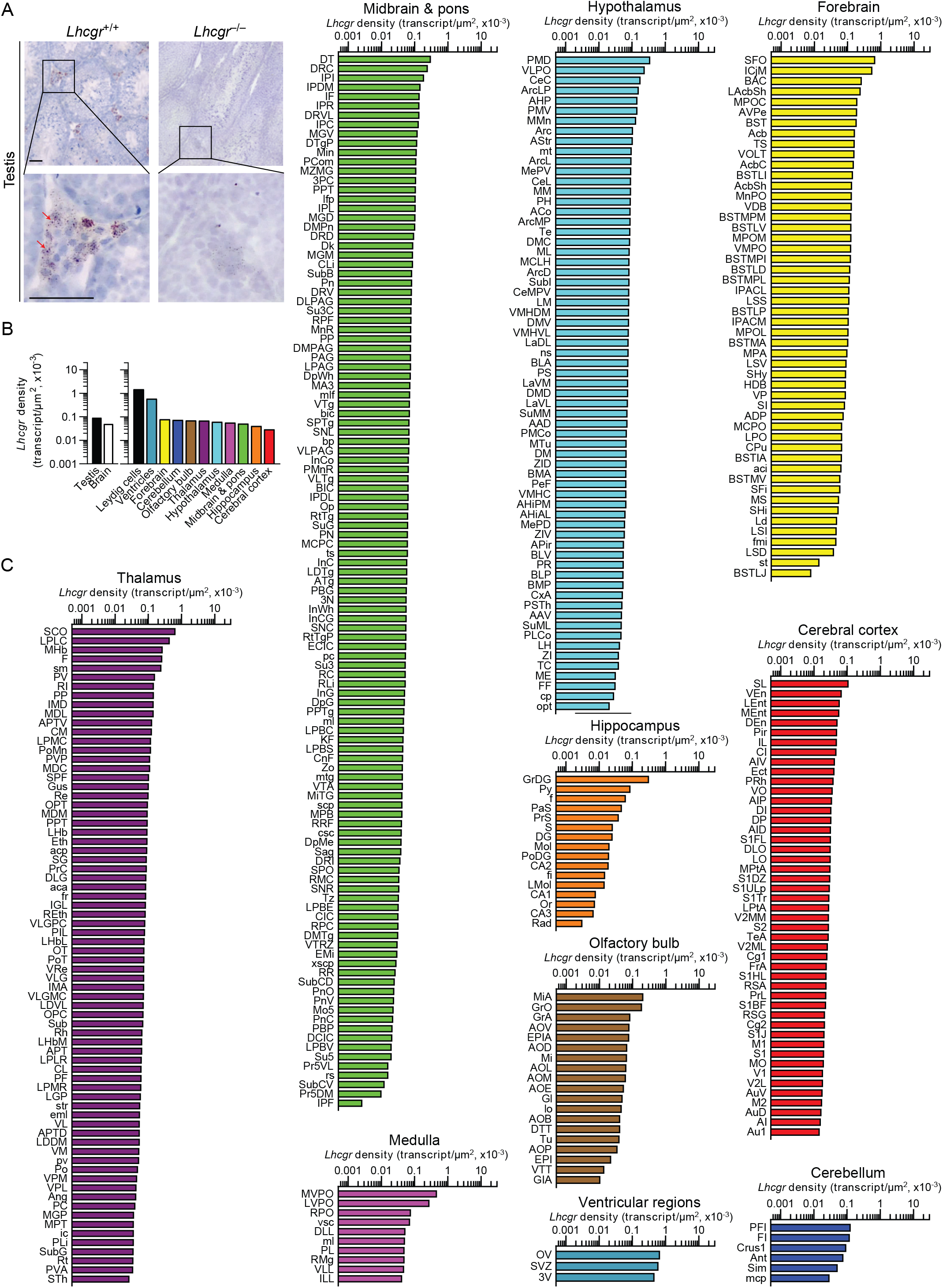
*Lhcgr* Expression in the Mouse Brain. **(A)** RNAscope signals were detected in the Leydig cells, but not juxtaposed Sertoli cells, in the mouse testis, confirming probe specificity. Scale bar: 25 μm. **(B)** *Lhcgr* transcript density in the testis and various brain regions detected by RNAscope. **(C)** *Lhcgr* transcript density in nuclei and sub-nuclei of the ventricular regions, forebrain, cerebellum, olfactory bulb, thalamus, hypothalamus, medulla, midbrain & pons, hippocampus and cerebral cortex.

We recently reported the expression of FSHRs in mouse, rat and human brains, particularly in AD–vulnerable regions, including hippocampus and cortex (30). We also found that FSH exacerbated AD–like neuropathology and cognitive decline in *3xTg, APP/PS1* and *APP*-KI mice, while the inhibition of FSH action rescued this phenotype. Most notably, shRNA– mediated knockdown of the *Fshr* in the hippocampus prevented the onset of AD–like features (30). Here, using RNAscope, we report the expression of *Fshrs* at the single–transcript resolution in 353 brain nuclei and sub–nuclei—and suggest that FSHRs in the brain may have roles beyond cognition. Probe specificity was established by a positive signal in testicular Sertoli cells, and an absent signal in juxtaposed Leydig cells and in the testes of *Fshr^−/−^* mice— as negative controls (Fig. 3A). Immunofluorescence confirmed the expression of FSHRs in NeuN–positive neurons, but not in GFAP–positive glial cells or IBA1–positive microglia (Fig. 3B).

**Figure 3:**
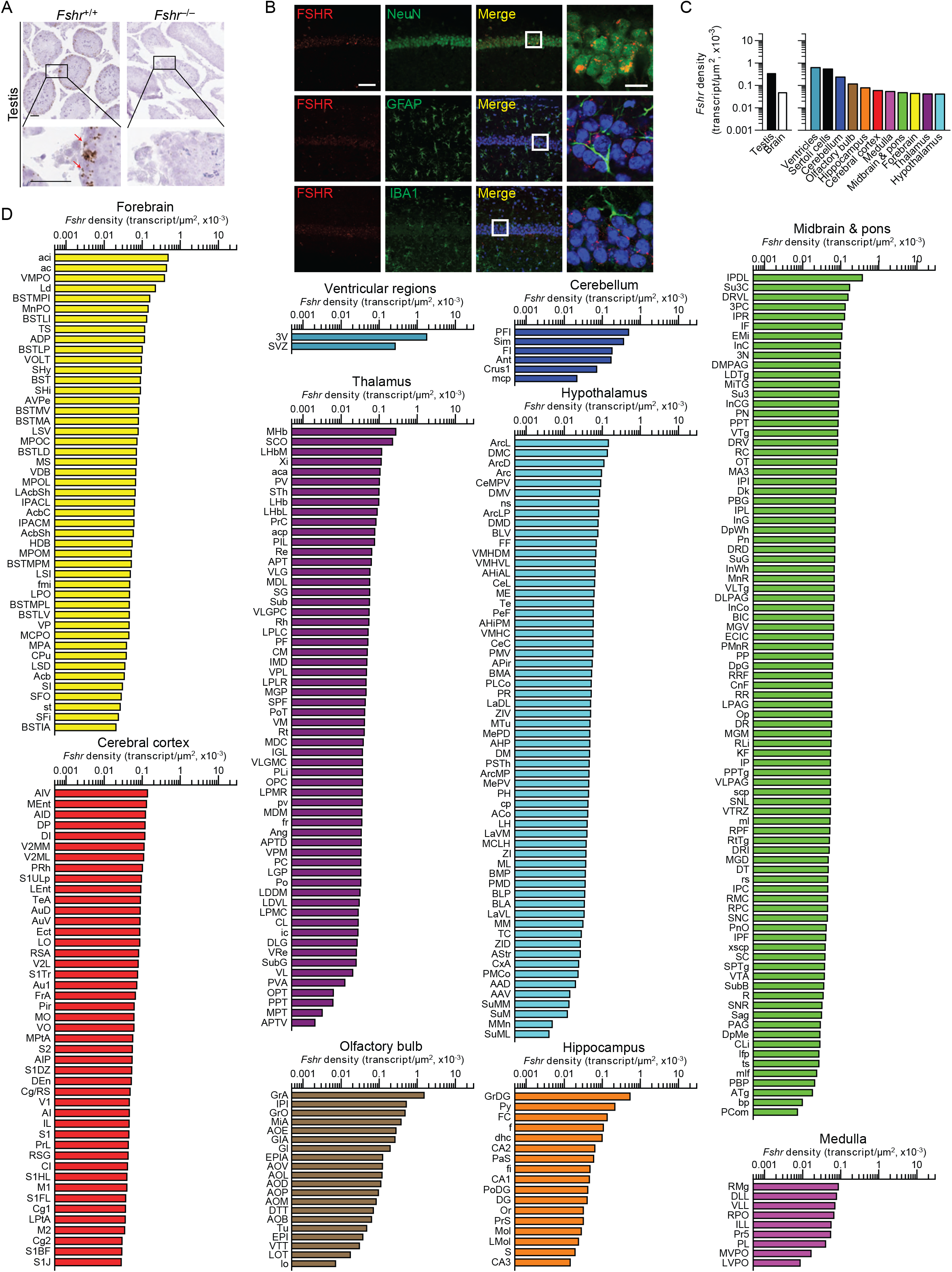
*Fshr* Expression in the Mouse Brain. **(A)** RNAscope signals were detected in the Sertoli cells, but not juxtaposed Leydig cells, in the mouse testis, confirming probe specificity. Scale bar: 50 μm. **(B)** FSHR immunofluorescence (red) was colocalized with NeuN–positive neurons, but not with GFAP–positive glial cells or IBA1–positive microglia. Scale bar: 100 μm (magnified view, 10 μm). **(C)** *Fshr* transcript density in the testis and various brain regions detected by RNAscope. **(D)** *Fshr* transcript density in nuclei and sub-nuclei of the ventricular regions, cerebellum, olfactory bulb, hippocampus, cerebral cortex, medulla, midbrain & pons, forebrain, thalamus and hypothalamus.

*Fshr* transcript density was highest in the ventricular region, followed in descending order, by the cerebellum, olfactory bulb, hippocampus, cerebral cortex, medulla, midbrain and pons, forebrain, thalamus, and hypothalamus (Fig. 3C). Within each region, respectively, the highest transcript densities were as follows: ependymal layer of the third ventricle (slightly higher than the testicular Sertoli cells); PFI in the cerebellum; GrA in the olfactory bulb; GrDG in the hippocampus; AIV in the cerebral cortex; RMg in the medulla; MHb in the thalamus; IPDL in midbrain and pons; aci in the forebrain; and ArcL in the hypothalamus (Fig. 3D). Raw transcript counts in each region and representative micrographs are shown in Supplementary Figs. 5 and 6, respectively.

We used ViewRNA to examine the expression of *FSHR* transcripts in specific regions of the human brain (Fig. 4A). Expression was noted in neuronal cells co–expressing the non-coding RNA *MALAT1* in the GrDG—consistent with the RNAscope data in mouse brain—and in the parahippocampal cortex. This latter data is consistent with *FSHR* expression in a population of excitatory glutamatergic neurons noted in human brain by 10X single cell RNA–seq (Allen Brain Atlas). Affymetrix microarray analysis confirmed *FSHR* expression in the frontal, cingulate, temporal, parietal and occipital sub–regions of human cortex in postmortem normal and AD brains (Fig. 4B). Interestingly, *FSHR* expression trended to be higher in the frontal cortex of the AD brains compared to that of unaffected brains (*P*=0.060). In all, the data suggest that, beyond a primary role in regulating cognition, brain FSHRs may have a wider role in the central regulation.

**Figure 4:**
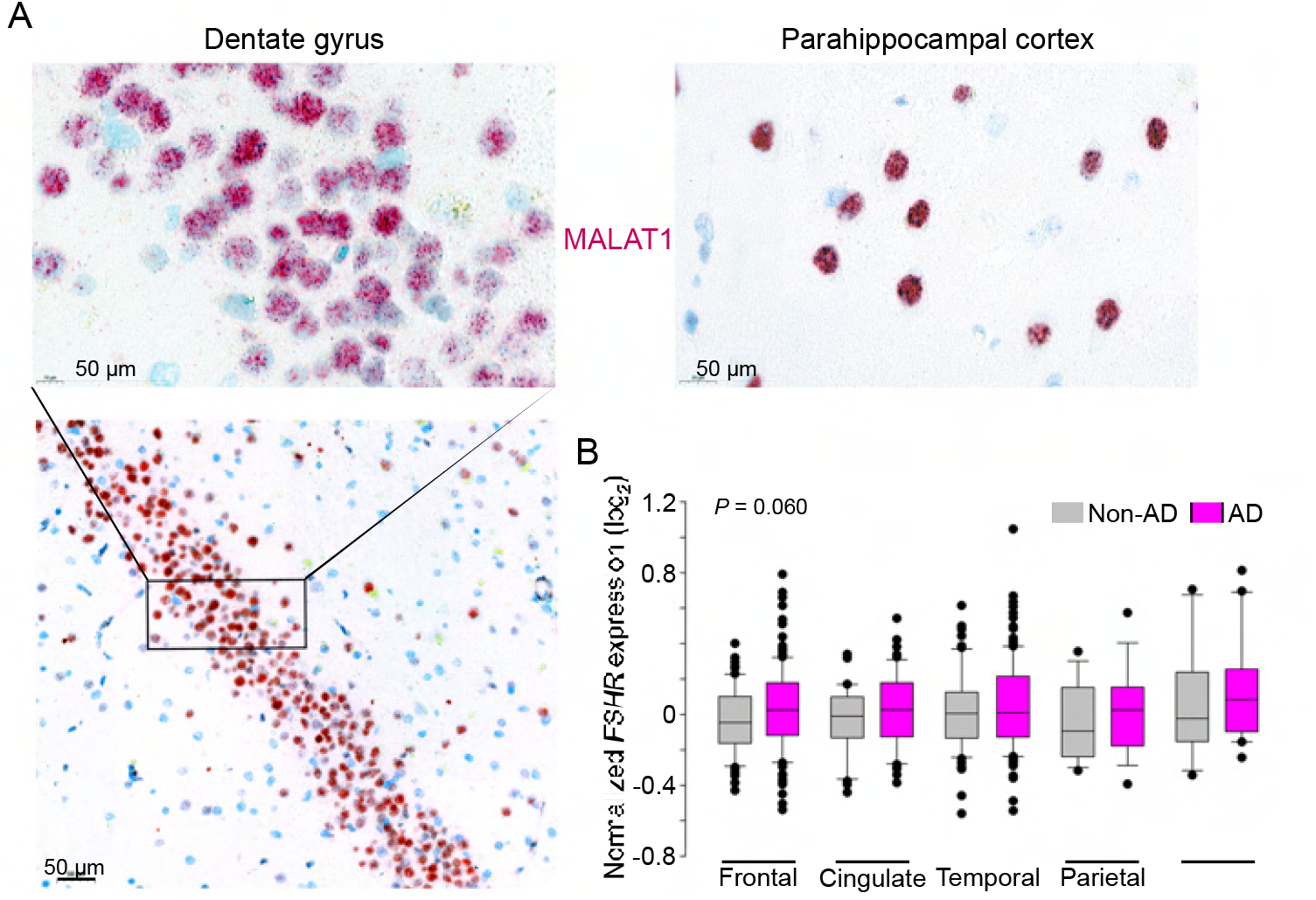
*FSHR* Expression in the Human Brain. **(A)** *FSHR* expression in the human hippocampus and parahippocampal cortex was detected by ViewRNA in neuronal cells that coexpress the non–coding RNA *MALAT1*. **(B)** *FSHR* mRNA expression in the frontal, cingulate, temporal, parietal and occipital sub–regions of human cortex in postmortem normal and AD brains (Affymetrix microarray, from GEO accession: GSE84422).

## DISCUSSION

The past decade has witnessed the unravelling of non–traditional physiologic actions of anterior pituitary glycoprotein hormones, and hence, the unmasking of functional receptors in bone, fat, brain, and immune cells, among other organs (1, 3, 4, 31–34). We report here for the first time that *Tshrs, Lhcgrs* and *Fshrs* are expressed in multiple brain regions. The data provide new insights into the distributed central neural network of anterior pituitary hormone receptors, particularly in relation to their role in regulating the somatic tissue function. Specifically, we find a surprising and striking overlap in central neural distribution of the three receptors—with highest transcript densities in the ventricular regions. Furthermore, at least for the TSHR and FSHR, expression levels in ependymal layer of the third ventricle was similar to that of the thyroid follicular cells and testicular Sertoli cells, respectively. *Albeit* intriguing, this may suggest a primary role for these receptors in central neural regulation.

Among 173 *Tshr*–positive brain regions, sub–regions and nuclei, the tanycyte–containing ependymal layer of the third ventricle displayed the highest *Tshr* transcript number and density. This region is juxtaposed to the anterior pituitary that produces TSH in response to hypothalamic thyrotropin–releasing hormone (TRH). Furthermore, TSH has been reported to be expressed in the hypothalamus (35, 36). It is therefore possible that a yet uncharacterized central TSH–TSHR feedback circuit may directly regulate the hypothalamic–pituitary–thyroid axis, thought solely to be controlled by thyroid hormones. To add to this complexity, thyroxine to triiodothyronine conversion occurs in tanycytes (37), which calls into question whether central TSH actions regulate thyroid hormone metabolism in these cells, and/or directly modulate hypothalamic TRH neuronal projections.

The forebrain and olfactory bulb also displayed abundant *Tshr* transcripts, with the highest density in the nucleus of the horizontal limb of the diagonal band (HDB) of the forebrain and ventral tenia tecta (VTT) of the olfactory bulb. These regions are involved, respectively, in learning and odor processing (38–42). In the hypothalamus, the highest density was found in medial tuberal nucleus (MTu), which controls ingestive behaviors and metabolism (43). Finally, we found more recently that the modulation of TSHRs in the bed nucleus of the stria terminalis (BNST), which receives direct afferents from the MTu (44), influences anxiety responses, suggesting that TSHR signaling might, in fact, mediate psychosocial behaviors.

While LH has a key role in reproduction and sexual development, we found 401 brain regions, sub–regions and nuclei expressing *Lhcgrs*. There were nominal differences in *Lhcgr* expression in many brain regions, but the ventricles stood out as having the highest *Lhcgr* density. Two regions deserve special mention. The *Lhcgr–rich* mitral cell layer of the accessory olfactory bulb (MiA) has a known role in scent communication during mating (45–48). A growing body of evidence suggests that men are attracted to cues of impending ovulation in women, raising an intriguing question on whether cycling hormones affect men’s attraction and sexual behavior (45, 48). The broader question is whether LH surges in women during cycling may, in fact, alter male sexual behavior through central mechanisms. Second, a high *Lhcgr* density in the subfornical organ (SFO) of the forebrain was surprising. SFO sends efferent projections to the organum vasculosum of the lamina terminalis (OVLT) (49, 50), which is surrounded by GnRH neurons and contains estrogen receptors (ESRs) (51). We therefore speculate that circumventricular interactions between LHCGRs, LH, GnRH, and ESR underpin the central regulation of reproduction.

RNAscope revealed 353 *Fshr*–expressing brain regions, sub–regions and nuclei. Highest expression was noted in the tanycyte–rich ependymal layer, not surprisingly given its anatomical proximity to the anterior pituitary gland where FSH is produced in response to hypothalamic gonadotropin–releasing hormone (GnRH). The functional significance of *Fshr* s expressed in the cerebellum, particularly in the paraflocculus (PFI), is yet unknown. However, other *Fshr*–high sub–regions, including the granular cell layer of the accessory olfactory bulb (GrA), granular layer of the dentate gyrus (GrDG) and agranular insular cortex (AIV), have known associations with odor processing, learning, memory formation and anticipation of reward (52–54). It is possible that the anosmia of Kallman syndrome, with unclear etiology, may arise from a dysfunctional FSHR–olfaction circuitry. We also find that inactivation of the hippocampal *Fshr* blunts the cognitive impairment and AD–like neuropathology induced by ovariectomy in *3xTg* mice. This data, together with gain– and loss–of–function studies suggest that hippocampal and cortical FSHRs could represent therapeutic targets for AD.

In all, our results provide compelling evidence for multiple central nodes being targets of the anterior pituitary glycoprotein hormones—a paradigm–shift that does not conform with the dogma that pituitary hormones are solely master regulators of single bodily processes. Through the intercession of emerging technologies, we compiled the most complete atlas of glycoprotein hormone receptor distribution in the brain at a single–transcript resolution. In addition, we have identified brain sites with the highest transcript expression and density, findings that are imperative towards a better understanding of the neuroanatomical and functional basis of pituitary hormone signaling in the brain. This understanding should provide the foundation for innovative pharmacological interventions for a range of human diseases, wherein direct actions of pituitary hormones, have been implicated, importantly, Alzheimer’s disease.

## METHODS

### Mice

We used *Tshr^+/-^* (strain #004858, Jackson Laboratory), *Lhcgr^-/-^* (strain #027102, Jackson Laboratory), *Fshr^-/-^* mice (55) and their wild type littermates in this study. Adult male mice (~3 to 4–month–old) were housed in a 12 h:12 h light:dark cycle at 22 ± 2 °C with *ad libitum* access to water and regular chow. All procedures were approved by the Mount Sinai Institutional Animal Care and Use Committee (approval number IACUC-2018-0047) and are in accordance with Public Health Service and United States Department of Agriculture guidelines.

### RNAscope

Mouse brain tissue was collected for RNAscope. Briefly, mice were anesthetized with isoflurane (2 to 3 % in oxygen; Baxter Healthcare, Deerfield, IL) and transcardially perfused with 0.9% heparinized saline followed by 4% paraformaldehyde (PFA). Brains were extracted and post–fixed in 4 % PFA for 24 hours, dehydrated and embedded into paraffin. Coronal sections were cut at 5 μm, with every tenth section mounted onto ~20 slides with 2–6 sections on each slide. This method allowed to cover the entire brain and to eliminate the likelihood of counting the same transcript twice. Sections were air dried overnight at RT and stored at 4 °C until required.

Simultaneous detection of mouse *Tshr, Lhcgr* and *Fshr* was performed on paraffin sections using RNAscope 2.5 LS Multiplex Reagent Kit and RNAscope 2.5 LS Probes, namely Mm-TSHR, Mm-LHCGR and Mm-FSHR (Advanced Cell Diagnostics, ACD). RNAscope assays on thyroid glands and testes (positive controls for *Tshr* and *Lhcgr/Fshr*, respectively), as well as brains from knockout mice (negative controls), were performed in parallel.

Slides were baked at 60 °C for 1 hour, deparaffinized, incubated with hydrogen peroxide for 10 minutes at room temperature, pretreated with Target Retrieval Reagent for 20 minutes at 100 °C and with Protease III for 30 minutes at 40 °C. Probe hybridization and signal amplification were performed *per* manufacturer’s instructions for chromogenic assays.

Following RNAscope assay, the slides were scanned at 20x magnification and the digital image analysis was successfully validated using the CaseViewer 2.4 (3DHISTECH, Budapest, Hungary) software. The same software was employed to capture and prepare images for the figures in the manuscript. Detection of *Tshr*–, *Lhcgr*– and *Fshr*–positive cells were also performed using the QuPath-0.2.3 (University of Edinburgh, UK) software based on receptor intensity thresholds, size and shape.

### Histology and Immunofluorescence

Heterozygous *Tshr^+/-^* mice in which a GFP cassette replaced exon 1 of the *Tshr* gene and their *Tshr^+/+^* littermates were euthanized with carbon dioxide and perfused transcardially with 0.9 % heparinized saline followed by 4 % PFA in 0.1 M phosphate buffered saline (PBS; pH 7.4). Brains were collected and post–fixed in the same fixative for overnight at 4 °C, then transferred to a 30 % sucrose solution in 0.1 M PBS with 0.1 % sodium azide and stored at 4 °C until they were sectioned on a freezing stage sliding microtome at 30 μm. Sections were stored in 0.1 M PBS solution with 0.1 % sodium azide until processed for double immunofluorescence.

For the double-label fluorescent immunohistochemistry, free–floating brain sections were rinsed in 0.1 M PBS (2 × 15 min) followed by a 30 min blocking in 3 % normal horse serum (Vector Laboratories, Burlingame, CA) and 0.3 % Triton X-100 in 0.1 M PBS. Sections were incubated with a mixture of primary rabbit anti–GFP antibody (1:500; catalog #SP3005P, OriGene, Rockville, MD) and mouse anti–NeuN antibody (1:1000; catalog #ab104224, Abcam, Cambridge, MA) for 18 h. Sections were then incubated with the secondary donkey anti–rabbit Alexa 488 (1:700; catalog #711-545-152, Jackson Immunoresearch, West Grove, PA) and donkey anti–mouse DyLight 594 (1:700; catalog #DK-2594, Vector Laboratories, Burlingame, CA) antibodies in 0.1 M PBS for 3 hours at room temperature. For immunohistochemical controls, the primary antibody was either omitted or pre–adsorbed with the immunizing peptide overnight at 4 °C resulting in no immunoreactive staining. In addition, we expectedly did not detect GFP immunoreactivity (-ir) in the *Tshr^+/+^* littermates, as the *Tshr* gene was intact and did not express GFP. Sections were mounted onto slides (Superfrost Plus) and cover–slipped using ProLong Gold Antifade Reagent (Life Technologies, Grand Island, NY). All steps were performed at room temperature.

For immunofluorescence staining for FSHR, free-floating brain sections were incubated overnight at 4°C with primary anti-FSHR (1:200; catalog # PA5-50963, ThermoFisher), anti-NeuN (1:300; catalog #MAB377, Sigma-Aldrich), anti-GFAP (1:400; catalog #MAB360, Sigma-Aldrich) or anti-IBA1 (1:500; catalog # PA5-18039, ThermoFisher) antibodies. After washing with Tris-buffered saline, the sections were incubated with a mixture of labelled secondary antibodies for detection. DAPI (Sigma-Aldrich) was used for staining nuclei.

### Microarray Analysis

Affymetrix Human Genome U133 Plus 2.0 Array data for *FSHR* expression in the frontal, cingulate, temporal, parietal and occipital cortex from both AD and non-AD human brains were curated from a previously published dataset (GEO accession #GSE84422) (56).

### Quantitative PCR

For quantitative RT-PCR performed on homogenates of brain tissues, total RNA from the hypothalamus and the hippocampus isolated from five *Tshr^+/+^* mice was extracted using an RNeasy Mini kit (Qiagen) *per* manufacturer’s protocol. Thyroid and the liver tissues were used as positive and negative controls, respectively. RNA was treated with DNAse I (Invitrogen), and reverse transcribed using the SuperScript II Reverse Transcriptase (Thermo Fisher Scientific). qPCR was performed with a QuantStudio 7 Real-Time PCR system (Applied Biosystems). PCR reaction mix consisted of first–strand cDNA template, exon-spanning primer pairs, and SYBR Green PCR master mix (Thermo Fisher Scientific). Expression of the selected targets was compared to that of a panel of normalizing genes *(Rps11, Tubg1* and *Gapdh)* measured on the same sample in parallel on the same plate, giving a Ct difference (ΔCt) for the normalizing gene minus the test gene. Relative expression levels were calculated by 2^-ΔΔCt^ using thyroid as the reference tissue.

### Quantitation, Validation and Statistical Analysis

Immunofluorescent images were viewed and captured using 10x or 20x objectives with an Observer.Z1 fluorescence microscope (Carl Zeiss, Germany) with appropriate filters for Alexa 488, Cy3 and DAPI. The captured GFP and NeuN images were evaluated and overlaid using AxioVision v.4.8 software (Carl Zeiss, Germany) and ImageJ (NIH, Bethesda, MD).

Data were analyzed by two-tailed Student’s *t*-test using Prism v.9.3.1 (Graphpad, San Diego, CA). Significance was set at *P* < 0.05.

## Acknowledgements

Work at Icahn School of Medicine at Mount Sinai was performed at the Center for Translational Medicine and Pharmacology and was supported by U19 AG060917 to M.Z. and C.J.R.; R01 DK113627 to M.Z. and T.F.D.; R01 AG074092 and U01 AG073148 to T.Y and M.Z; and R01 AG071870 to S.-M.K., T.Y. and M.Z. M.Z. also thanks the Harrington Discovery Institute for the Innovator–Scholar Award. C.J.R. acknowledges support from the NIH (P20 GM121301).

## DATA AVAILABILITY

All data generated or analyzed during this study are included in the manuscript and supporting files.

---

**<u>Table 1:</u> Known Functions of TSHR, FSHR and LHCGR in the Brain.**

**Supplementary Figure 1:**
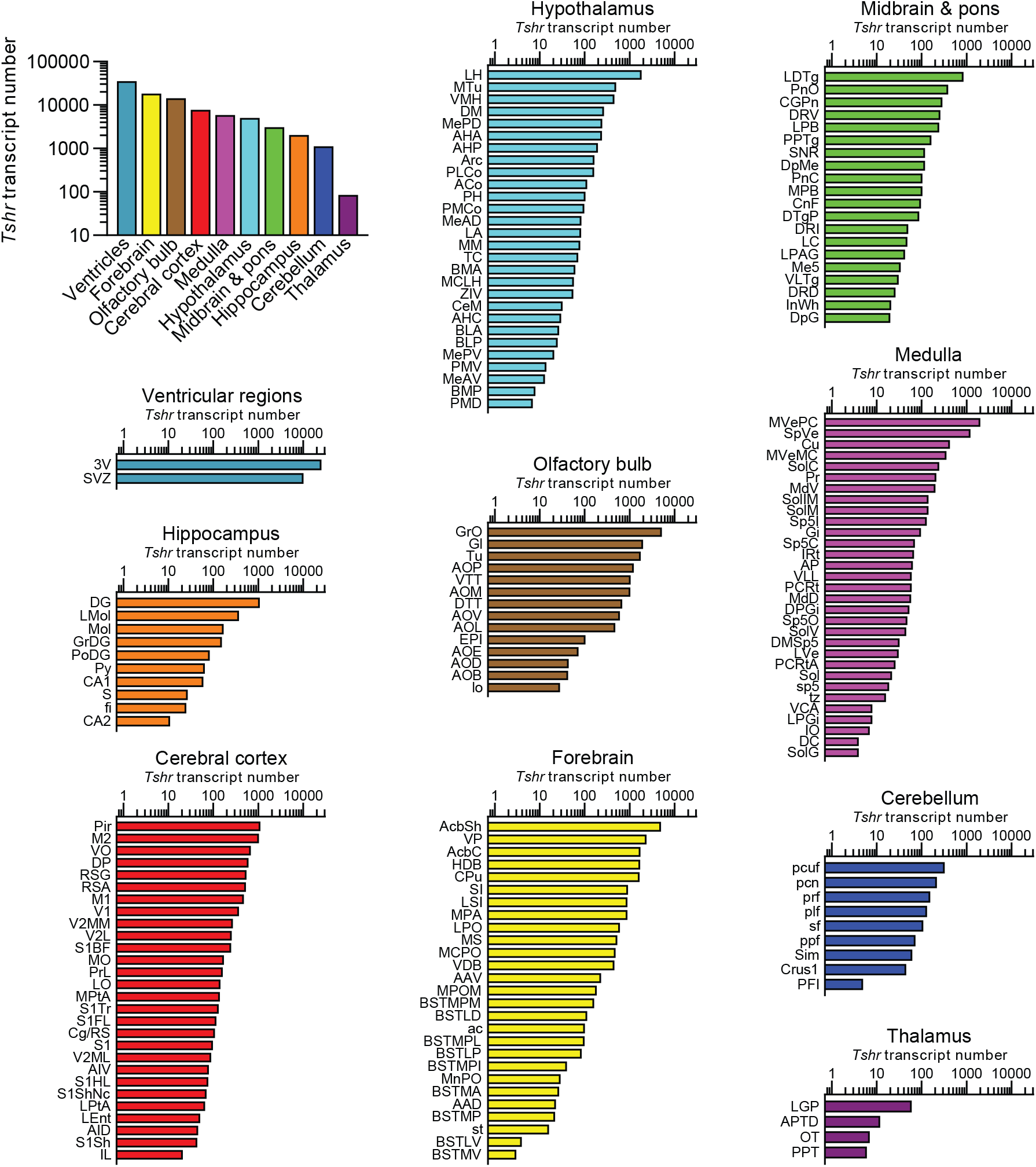
Raw *Tshr* transcript counts in each brain region, nuclei and subnuclei.

**Supplementary Figure 2:**
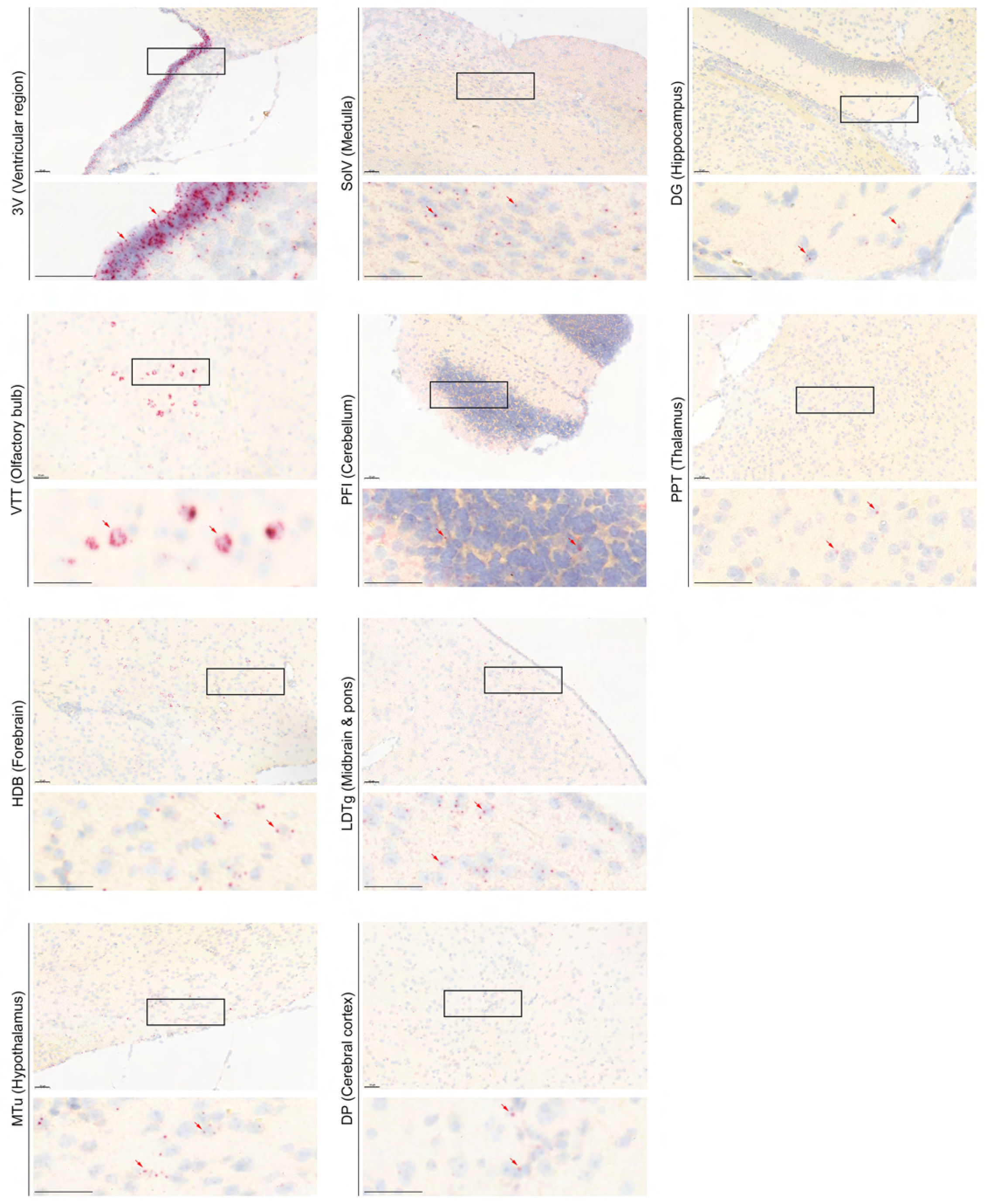
Representative RNAscope micrographs showing *Tshr* transcripts in the ependymal layer of the third ventricle (3V), ventral tenia tecta (VTT) of the olfactory bulb, nucleus of the horizontal limb of the diagonal band (HDB) of the forebrain, medial tuberal nucleus (MTu) of the hypothalamus, solitary nucleus, ventral part (SolV) of the medulla, paraflocculus (PFI) of the cerebellum, laterodorsal tegmental nucleus (LDTg) of the midbrain and pons, dorsal peduncular cortex (DP) of the cerebral cortex, dentate gyrus (DG) of the hippocampus, and posterior pretectal nucleus (PPT) of the thalamus. Scale bar: 50 μm.

**Supplementary Figure 3:**
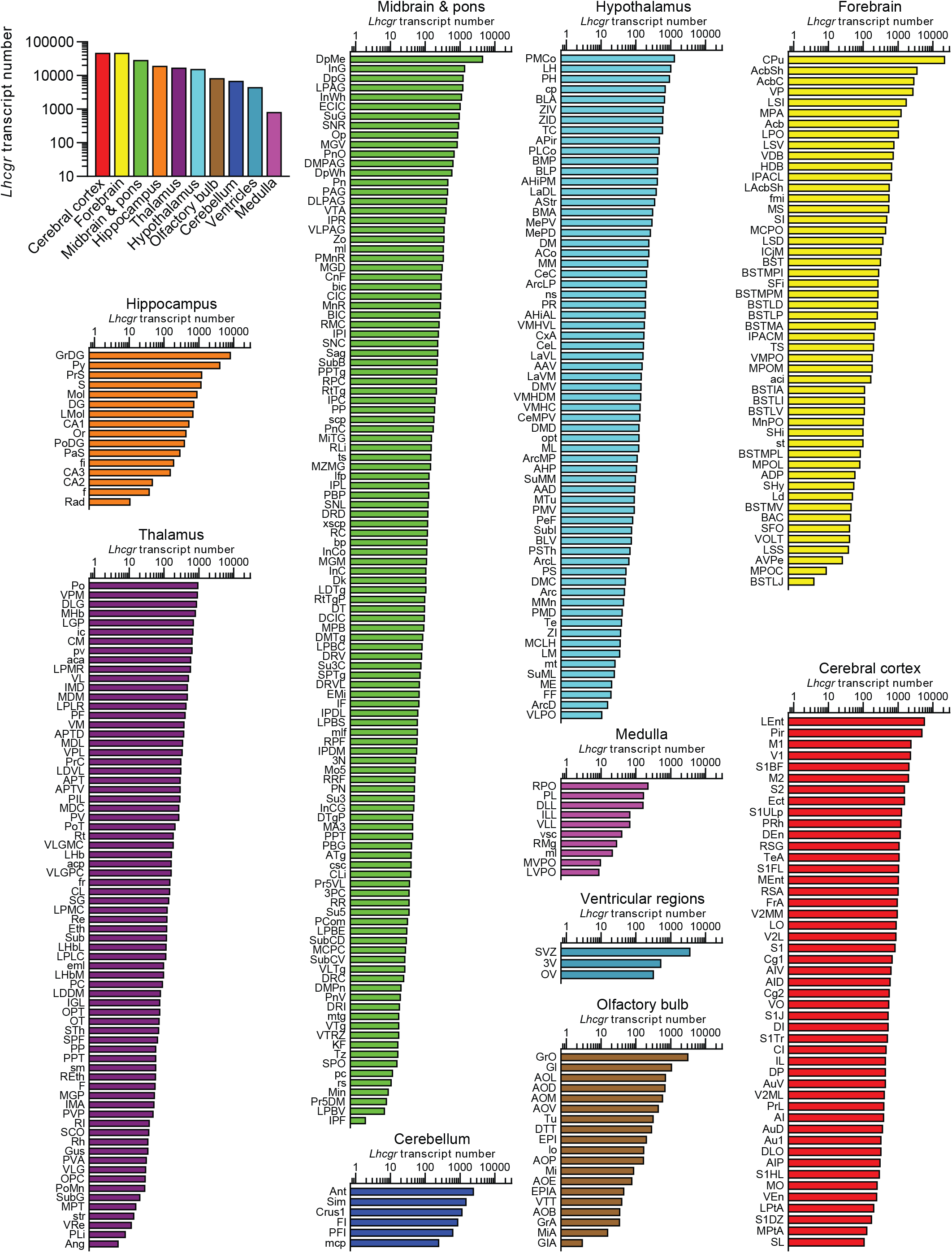
Raw *Lhcgr* transcript counts in each brain region, nuclei and subnuclei.

**Supplementary Figure 4:**
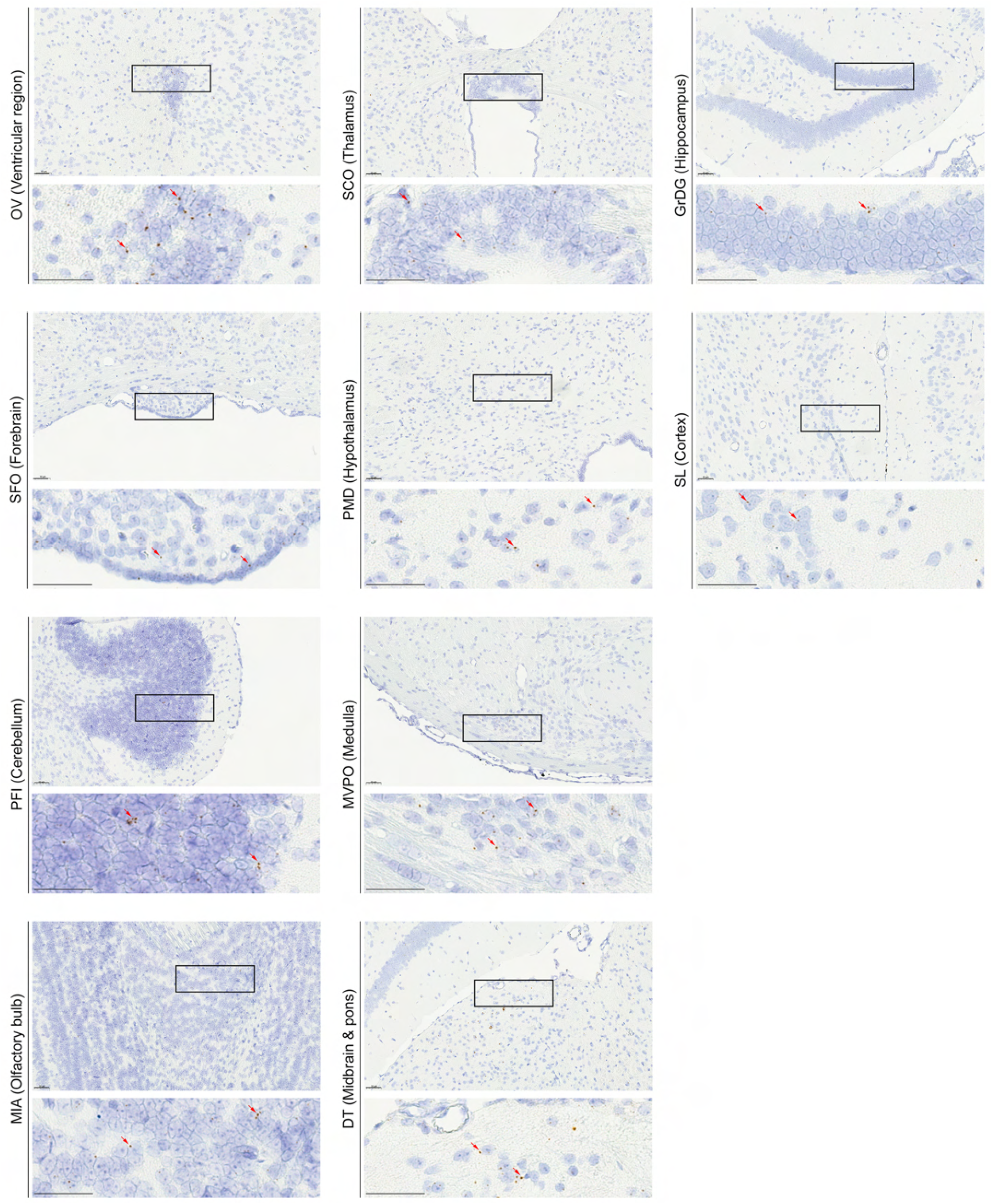
Representative RNAscope micrographs showing *Lhcgr* transcripts in the olfactory ventricle (OV), subfornical organ (SFO) of the forebrain, paraflocculus (PFI) of the cerebellum, mitral cell layer of the accessory olfactory bulb (MiA), subcommissural organ (SCO) of the thalamus, premammillary nucleus, dorsal part (PMD) of the hypothalamus, medioventral periolivary nucleus (MVPO) of the medulla, dorsal terminal nucleus of the accessory optic tract (DT) of the midbrain and pons, granular layer of the dentate gyrus (GrDG) of the hippocampus, and semilunar nucleus (SL) of the cerebral cortex. Scale bar: 50 μm.

**Supplementary Figure 5:**
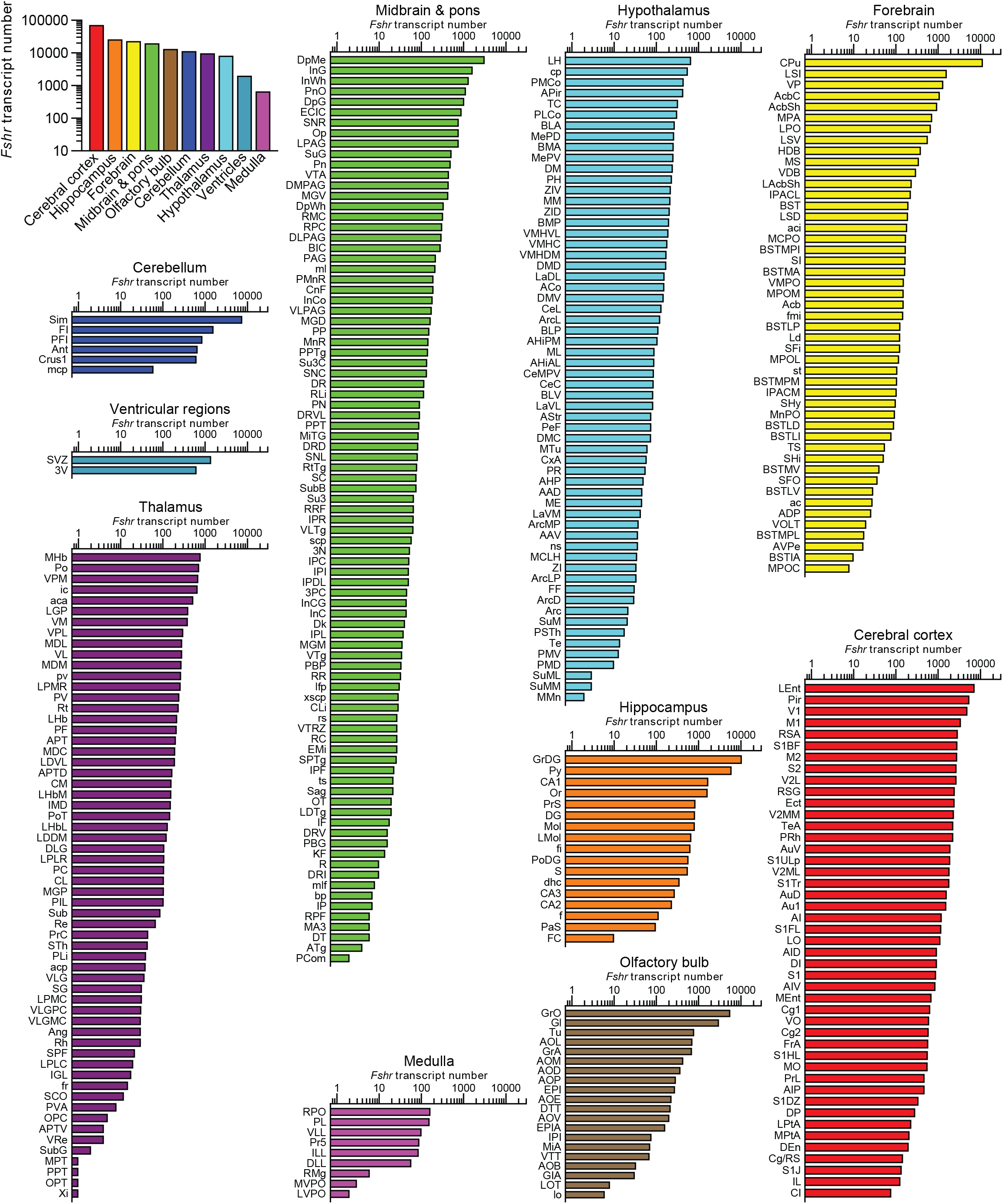
Raw *Fshr* transcript counts in each brain region, nuclei and subnuclei.

**Supplementary Figure 6:**
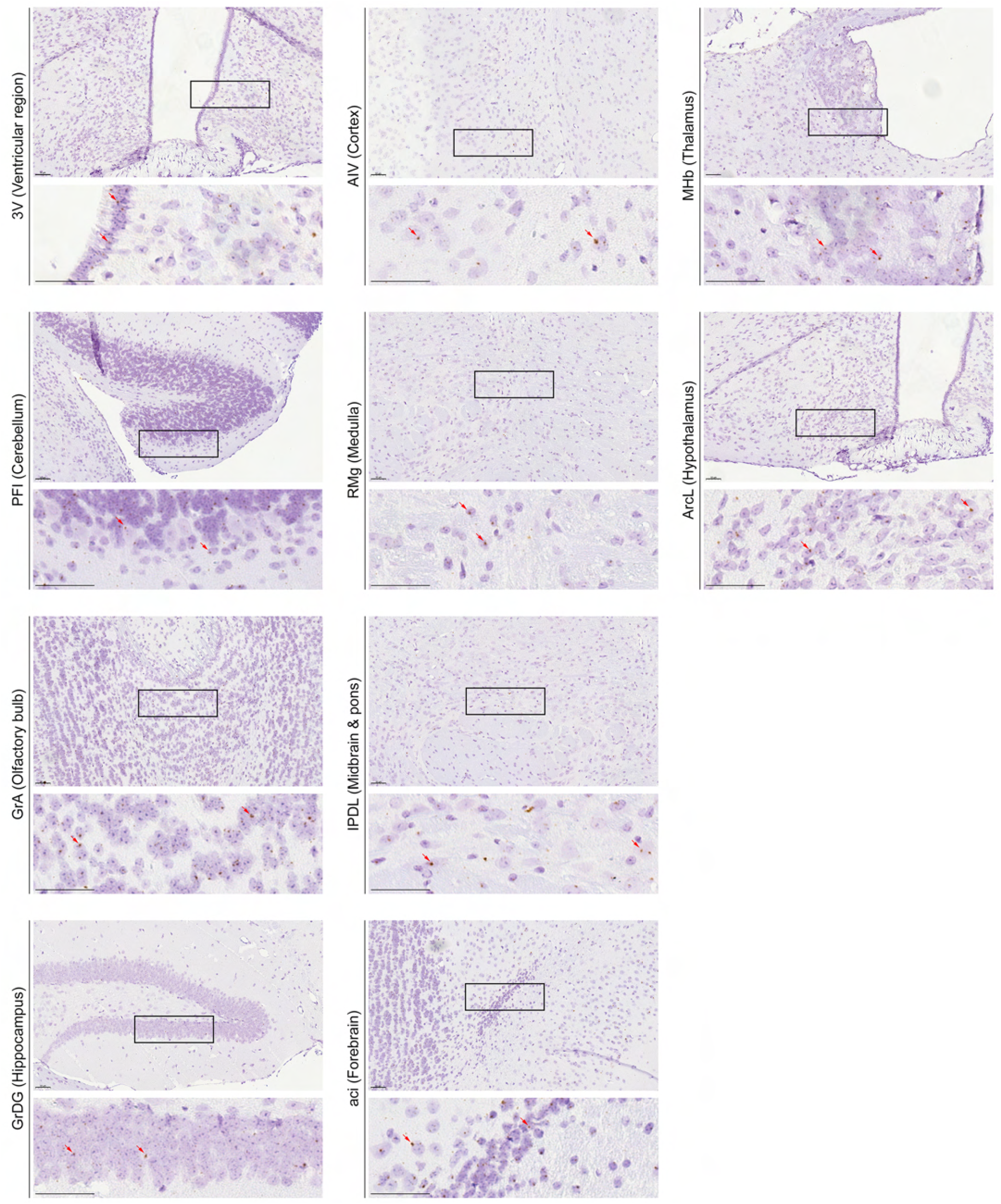
Representative RNAscope micrographs showing *Fshr* transcripts in the ependymal layer of the third ventricle (3V), paraflocculus (PFI) of the cerebellum, granule cell layer of the accessory olfactory bulb (GrA), granular layer of the dentate gyrus (GrDG) of the hippocampus, agranular insular cortex, ventral part (AIV) of the cerebral cortex, raphe magnus nucleus (RMg) of the medulla, interpeduncular nucleus, dorsolateral subnucleus (IPDL) of the midbrain and pons, anterior commissure, intrabulbar part (aci) of the forebrain, medial habenular nucleus (MHb) of the thalamus, and arcuate hypothalamic nucleus, lateral part (ArcL) of the hypothalamus. Scale bar: 50 μm.

## Appendix

Glossary of the brain regions, nuclei and subnuclei.

